# Selection-free, high frequency genome editing by homologous recombination of human pluripotent stem cells using Cas9 RNP and AAV6

**DOI:** 10.1101/252163

**Authors:** Renata M. Martin, Kazuya Ikeda, Nobuko Uchida, Kyle Cromer, Toshi Nishimura, Daniel P. Dever, Joab Camarena, Rasmus Bak, Anders Laustsen, Martin R. Jakobsen, Volker Wiebking, Vittorio Sebastiano, Hiromitsu Nakauchi, Matthew Porteus

## Abstract

Combination of genome editing and human pluripotent stem cells (hPSCs) offers a platform for *in vitro* disease modeling, drug discovery and personalized stem cell therapeutics. However, incorporation of large modifications using CRISPR/Cas9-based genome editing in hPSCs typically requires the use of selection markers due to low editing efficiencies. Here we report a novel editing technology in hPSCs using Cas9 protein complexed with chemically modified single guide RNA (sgRNA) and recombinant AAV6 (rAAV6) vectors for donor delivery without marker selection. With these components, we demonstrate targeted integration of a 2.2 kb DNA expression cassette in hPSCs at frequencies up to 94% and 67% at the *HBB* and *MYD88* loci, respectively. We used this protocol to correct the homozygous sickle cell disease (SCD) mutation in an iPSC line derived from a SCD patient with a frequency of 63%. This Cas9/AAV6 system allows for both the integration of large gene cassettes and the creation of single nucleotide changes in hPSCs at high frequencies, eliminating the need for multiple editing steps and marker selection, thus increasing the potential of editing human pluripotent cells for both research and translational applications.

## Main Text

Combining programmable nucleases such as ZFNs, TALENs and Cas9 with a donor template for DNA repair can allow precise genome editing in many cell types through homologous recombination (HR). The application of this genome editing technology to human pluripotent stem cells (hPSCs) has advanced disease modeling and regenerative medicine. Recently the CRISPR/Cas9 system, combined with delivery of a donor vector in the form of single-stranded oligodeoxynucleotides (ssODNs), has been applied to hPSCs to create small edits (<10bp) at high efficiency^1^.

This strategy enables the modeling of diseases caused by single or several base pair mutations, which is frequently the case for genetic disorders. The ssODN approach, however, does not allow the targeted integration of large gene cassettes which is needed for certain gene correction type approaches and for many research applications. For example, to make reporter cell lines to study a disease, targeted integration of large gene fragments (>1kb) is required in order to integrate a reporter gene as a marker for differentiation, which can be important for biological understanding and clinical application of hPSCs. Large gene integration is impractical using the current Cas9/ssODN technology due to inverse correlation between ssODN size and editing efficiency (most efficient size is 90bp)^2^. Other systems, such as iCRISPR, can yield high frequencies in embryonic stem cells (ESCs) of around 30%^3^ for small base pair edits, however this efficiency is dramatically reduced to <1%^4^ when large insertions are introduced. To obtain a large population of edited cells with a protocol that yields such low editing frequencies requires the use of a selection marker, which could affect cell function, thereby limiting research and clinical relevance. To avoid the possible influence of a selection marker, piggyBac or Cre/loxP systems have been used to excise this marker. However, both of these approaches require an additional editing step and the use of piggyBac requires a nearby TTAA site^5^.

It was recently shown that large ssODN donor vectors are able to incorporate large modifications at high editing frequencies when delivered directly into zygotes by microinjection^6^. Though microinjection is impractical for editing large populations of hPSCs, we hypothesized that efficient delivery of donor vectors into the nucleus is a limiting aspect for editing frequency. Along these lines, we previously reported that the combination of adeno-associated virus (AAV), chemically-modified singe-stranded guide RNA (sgRNA), and Cas9 elicits high editing frequencies in human primary CD34+ hematopoietic stem cells, a cell type that has been refractory to previous genome editing protocols, suggesting that AAV vectors may efficiently deliver large donor constructs to the nucleus^7–9^ To test this hypothesis, we assessed the targeting frequency at the *HBB* locus of a large expression cassette (2.2kb; GFP driven by a UbC promoter) in human H9 ESCs using AAV6 as a donor delivery vehicle (Figure 1a). We electroporated cells with Cas9 ribonucleoprotein (RNP) complexed with a chemically-modified sgRNA^10^ targeted to exon 1 of the *HBB* gene, then immediately (<5 minutes) afterward transduced the electroporated cells with AAV6 donor vector. Our UbC-GFP donor expresses GFP episomally, therefore we assessed editing frequency at an average of 42.27% + 0.07% after non-integrated AAV6 episomal expression was diluted out as the population of cells expanded post-editing, a process that occurred within 4 days (Figure 1b). Using this approach, we observed a population of highly edited ESCs stably expressing GFP at an average frequency of 53.7% + 0.11% compared to plasmid-based vector (0.34% + 0.002%) at four days post-editing (Figure 1c,d). We then sought to determine whether this same approach could achieve high editing frequencies in iPSC lines at similar efficiencies. Indeed, when targeting the *HBB* locus with our UbC-GFP AAV6 donor vector in five different induced pluripotent stem cell (iPSC) lines we observed editing frequencies at an average of 51.28% ± 0.12% (Figure 1e).

**Figure 1.**
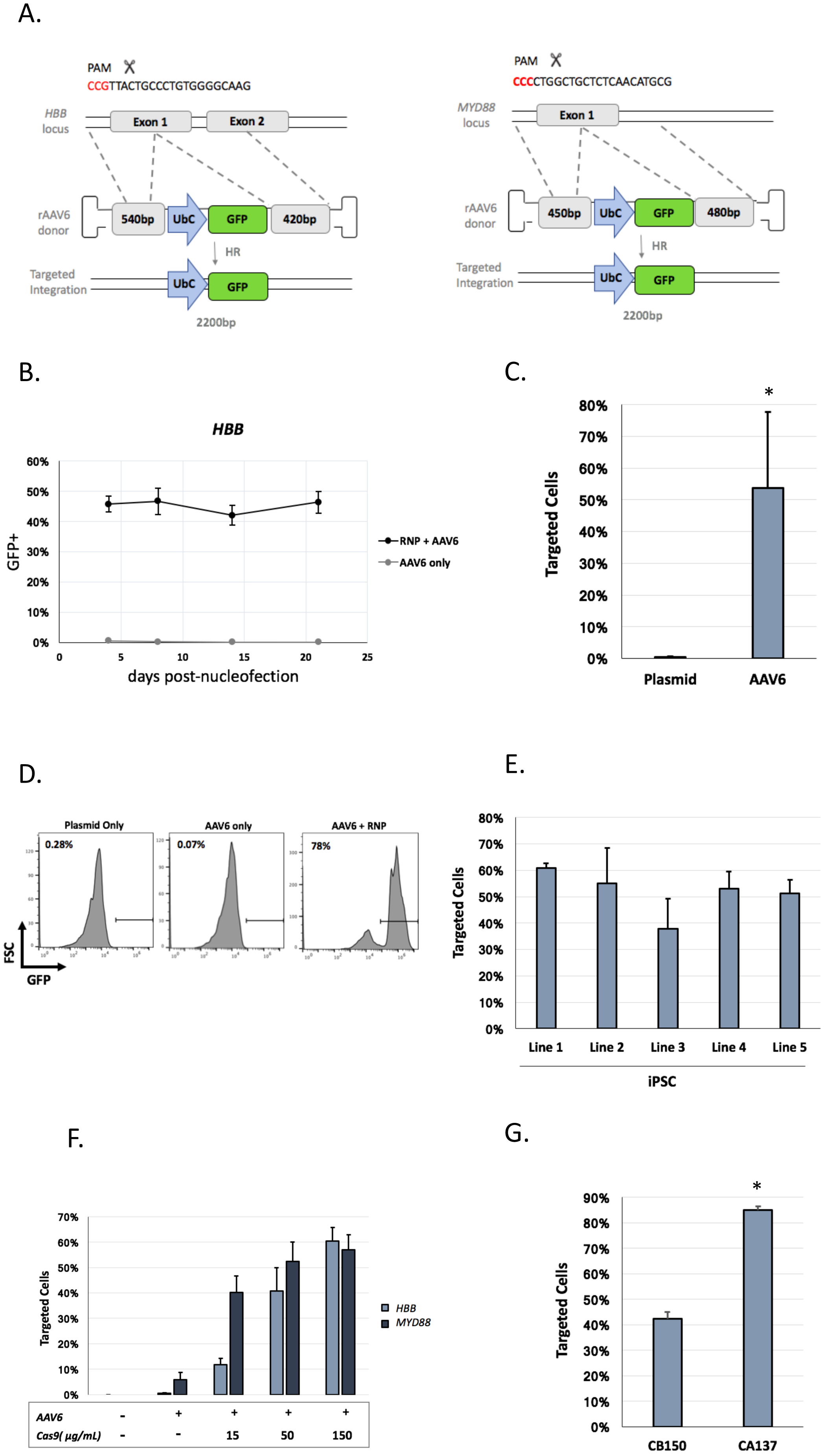
Optimization of Cas9/AAV6 editing in ESCs and iPSCs. **(a)** Schematic of targeted genome editing at the *HBB* (right) and *MYD88* (left) loci using CRISPR/Cas9 RNP and AAV6. Site-specific DSBs are created by Cas9 mainly between nucleotides 17 and 18 of the 20-bp target site, which is followed by the 5′-NGG-3′ PAM sequence (red). A DSB can be repaired by homologous recombination (HR) using a GFP-encoding rAAV6 homologous donor as a repair template. **(b)** Embryonic stem cells were edited with *HBB* donor and Cas9 RNP with CB-150 electroporation program. Data shows stable expression of GFP throughout the time course of 21 days (n=4). Editing frequencies as measured by GFP expression by flow cytometry at *HBB* 8 days post electroporation with CB-150 program of ES cells using either AAV6 or plasmid-based GFP donor delivery. Donor features are identical for the AAV6 and plasmid donor. AAV6 data is based on n=5 and plasmid data is based on n=3. p < 0.0005 based on unpaired T-test. **(d)** A representative FACS plot from a single replicate comparing plasmid vs. AAV6 + RNP. AAV6 only is shown as the control. **(e)** Genome editing at the *HBB* locus 8 days post nucleofection in 5 different iPSC lines using electroporation protocol CA-137. (n=2). No significant difference between cell line editing frequencies. **(f)** Gene editing frequencies 8 days post nucleofection at the *HBB* and *MYD88* loci using increasing Cas9/gRNA concentrations at a molar ratio of 1/3, respectively. (n=2 for *MYD88* and n=3 for *HBB* samples). **(g)** Comparison of the CB-150 and CA-137 electroporation programs for gene editing of ESCs with *HBB* Cas9 RNP followed by *HBB* AAV6 donor delivery. Percent GFP+ cells were quantified 8 days post-electroporation by flow cytometry. (n=4). If not mentioned otherwise, 150 μg/mL of Cas9 and CB-150 electroporation program were used. p < .00001 based(**c**) on unpaired T-test.

In an effort to further improve these targeting rates we next optimized variables within the protocol. Increasing the amounts of Cas9 during electroporation increased targeting frequencies at both the *HBB* and *MYD88* loci in a dose-dependent manner (Figure 1f). Finally, we tested different electroporation programs and found that CA137 protocol of the Lonza 4-D system was able to more than double the editing frequency at the *HBB* locus compared to the previously used CB150 (84% vs. 41%, respectively, Figure 1g).

Combining the fully optimized conditions (150μg/mL Cas9 with CA137 electroporation program), we observed 91% targeted integration at *HBB* and 59% at *MYD88* in ESCs (Figure 2a). To analyze whether cells possessed mono‐ or bi-allelic integrations, we performed single cell cloning of the two populations targeted at either *MYD88* and *HBB* by random colony picking. PCR analysis using primers annealing outside each homology arm determined that 17 out of 18 clones (94%) for *HBB* and 13 out of 21 clones (62%) for *MYD88* possessed at least one integration (Figure 2b and 2c). Editing occurred with such great frequency that we were even able to obtain a high percentage of bi-allelic integration events for *HBB* (74% of the screened clones). For *MYD88*, we obtained mostly mono-allelic clones (52%) and 10% bi-allelic clones (Figure 2c-d). We also performed off-target analysis at the two most probable off-target sites for both *HBB* and *MYD88* sgRNAs identified by the COSMID target prediction tool^7,10–13^. This analysis showed off-target activity for both sgRNAs with 16 out of 18 clones for *HBB* and 19 out of 21 clones for *MYD88* harboring off-target NHEJ events at one or both off-target sites (Supplemental figure 1a-d). Thus, high frequencies of on-target editing can be accompanied by high frequencies of INDELs, though the INDELs are occurring at sites of no known biologic significance.

**Figure 2.**
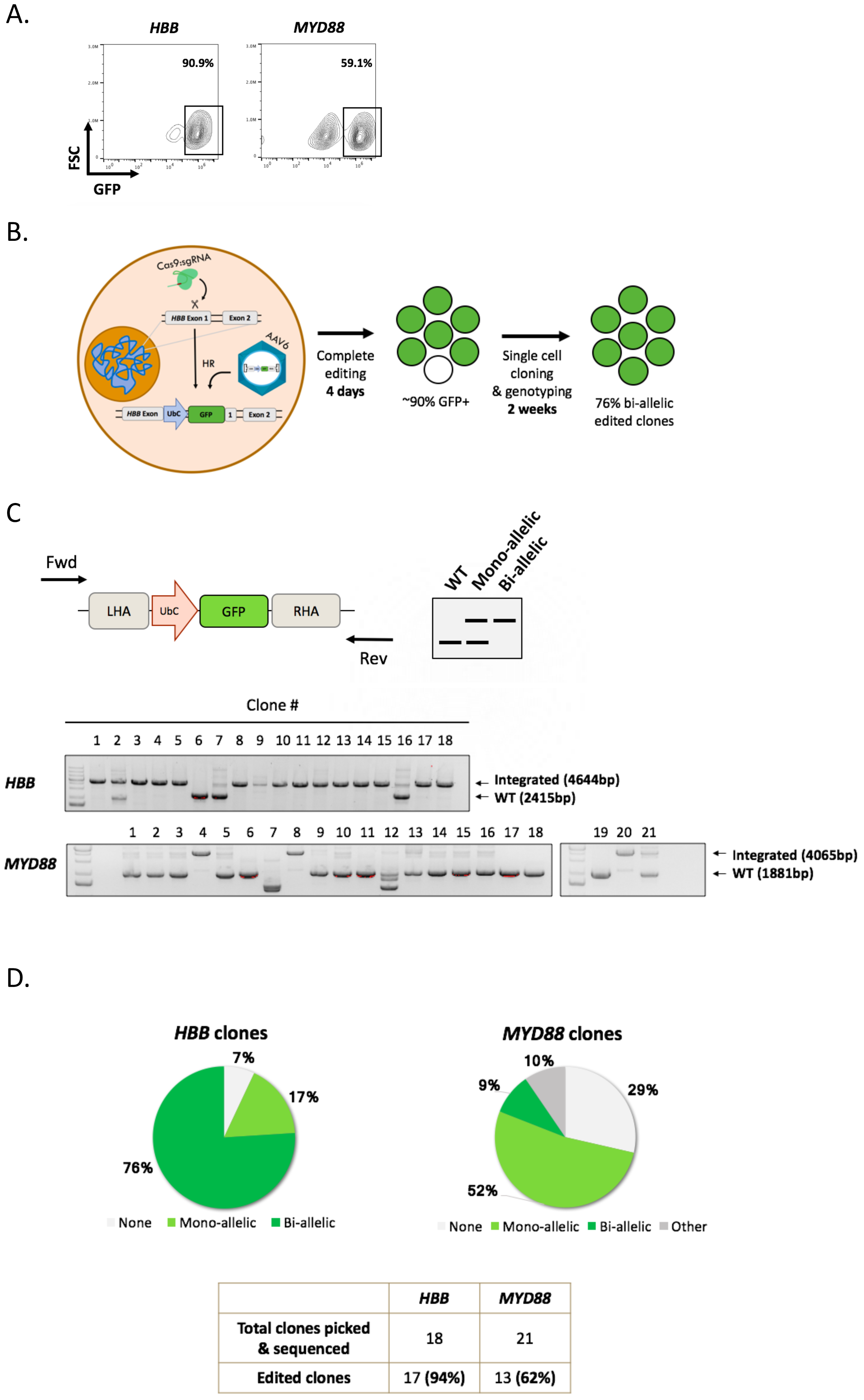
Analysis of AAV6 edited clones at *HBB* and *MYD88* loci. **(a)** Targeted integration at the *HBB* and *MYD88* loci using optimized conditions (150 μg/mL of Cas9 and CA-137 electroporation program). Editing frequencies were determined 8 days postelectroporation by flow cytometric analysis of GFP+ cells. **(b)** Schematic shows the gene editing process using Cas9 RNP and AAV6-mediated donor delivery followed by single cell cloning after four days which ultimately allows genotyping of clones. **(c)** Schematic depiction of primer locations for clonal genotyping of the *HBB* and *MYD88* loci with both primers binding the genomic regions outside the homology arms of the donor vectors (*top left*) and a schematic representation of an agarose gel with all three possible genotype PCR amplicons depicted (*top right*). Agarose gel images show genotypes of analyzed clones targeted at the *HBB* and *MYD88* loci, respectively (*bottom*) **(d)** Pie charts showing distribution of bi-allelic, mono-allelic, and WT clones from *HBB* and *MYD88*-targeted populations.

We next tested our system in a disease-relevant iPSC line homozygous for the Glu6Val sickle cell disease (SCD)-causing mutation^14^. Toward this end, we used a previously used AAV6 donor vector (Supplemental figure 2 that corrects the disease SNP (T→A) and introduces five silent mutations with sgRNA target sequence to prevent Cas9 from re-cutting and disrupting the gene^7^ (Figure 3a). Combining this SCD-correction donor and Cas9 RNP, we targeted the SCD iPSC line using the optimized protocol, and droplet digital PCR (ddPCR) analysis showed 63% allele correction (Figure 3b). As before, we confirmed these results using single cell cloning followed by sequencing, which confirmed the high correction frequency with 15 out of 22 corrected clones (68%; Figure 3c). Off-target analysis identified one corrected clone that did not contain off-target edits at the previous two off-target sites for the *HBB* sgRNA (clone #8; Figure 3d).

**Figure 3.**
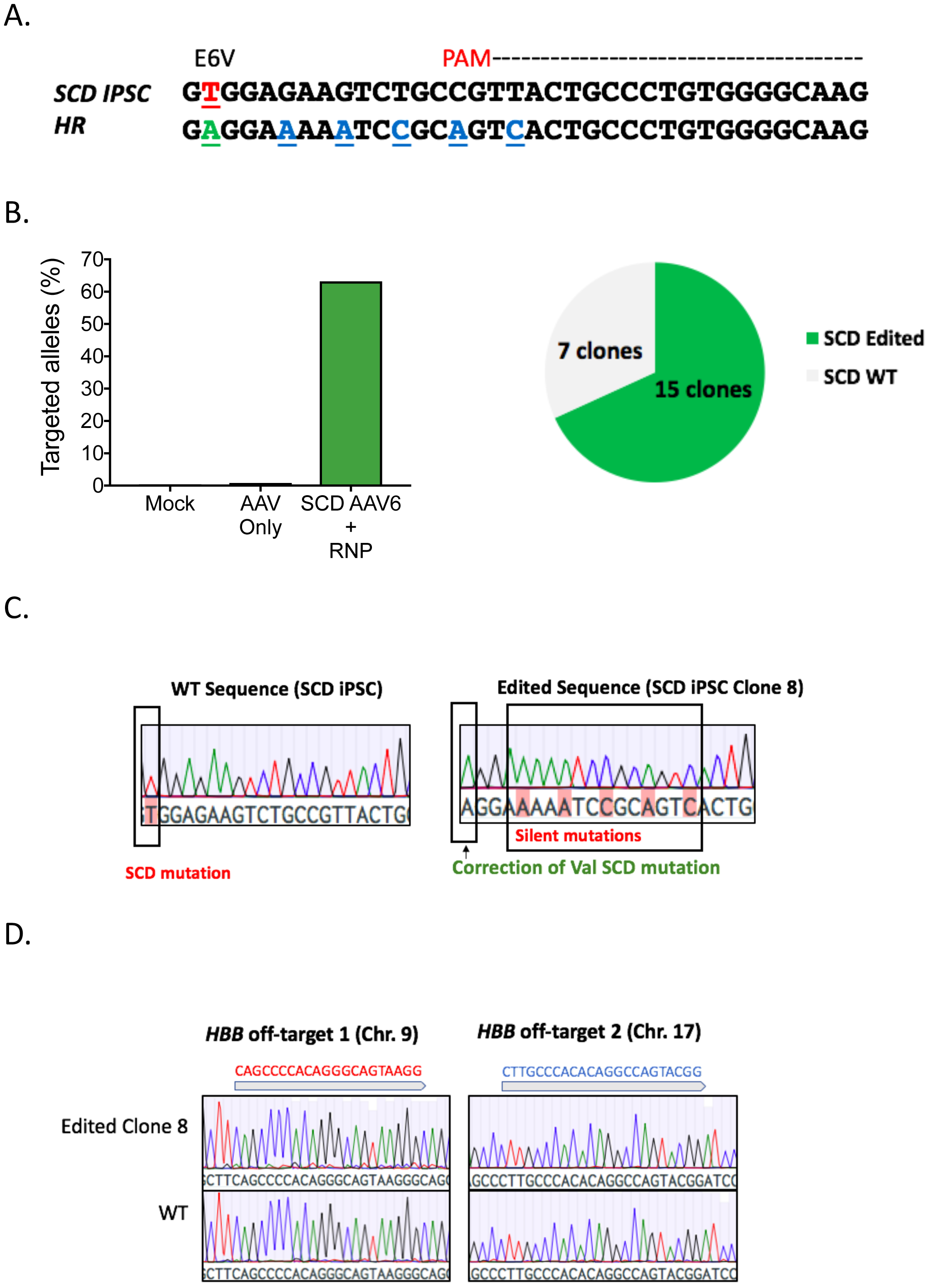
Sickle cell disease iPSC correction with AAV6 donor. **(a)** Sequence of the *HBB* and *MYD88* gene at the sickle cell disease-causing SNP with the *HBB* sgRNA target site depicted with dashes and PAM in red. Below is the sequence of the site after corrective HR. The corrective donor vector contains the WT sequence and silent mutations in the sgRNA target site to prevent Cas9 RNP from re-cutting the site. **(b)** Sickle cell disease iPSCs were edited with the corrective AAV6 donor and bulk genomic DNA samples were collected 8 days after electroporation. Droplet digital PCR (ddPCR) was used to measure targeted integration in the population (*left*) and sequencing of single cell clones was performed to confirmed the editing rates. **(c)** Sequence chromatograms of an unedited (*left*) and edited (*right*) SCD iPSC clone. The latter shows the correction of the sickle cell mutation and introduction of the intended silent mutations. **(d)** Sequence chromatograms from an unedited (*bottom*) and edited (*top*) SCD iPSC clone showing no off-target effects at two COSMID-predicted off-target sites.

In summary, our data confirms that Cas9 RNP and AAV6 is able to mediate targeted integration of large genes in ESCs and iPSCs at two different loci as well as precisely correct a disease-causing mutation in a patient-derived iPSC line, all at high frequencies. Importantly, these high editing frequencies enable population-based studies without the requirement for selection markers or large scale single cell cloning. Although clones were found to contain mutations at predicted off-target sites, none of these sites for either guide lie within or near genes are thus unlikely to elicit a significant cellular effect. However, this finding highlights the importance of careful sgRNAs selection. Specificity may be enhanced using engineered Cas9 variants with higher specificity^15^, truncated or chemically modified sgRNAs with lower affinity for off-targets^10^ or by careful titration of Cas9 RNP amounts. In fact, we did observe higher specificity when using the CB-150 electroporation program and with lower amounts of Cas9 (Supplemental Figure 1c-d). In addition to the highly improved editing frequencies in hPSCs using the Cas9/AAV6 system, we also demonstrated efficient integration of a large 2.2kb construct. This approach, using AAV6 to deliver donors up to 4.5kb, makes it possible to perform editing by cDNA knock-in in hPSCs as well, a strategy that would be critical for diseases with loss-of-function mutations scattered throughout a particular gene and would also be easily applied as a tool to generate genetically engineered human pluripotent cells for a variety of research purposes, such as to create stable transgenic lines or reporter lines.

In conclusion, we present a selection-free, one-step editing protocol that addresses insert size limitations and low editing efficiencies that have hindered the development of pluripotent stem cell technologies as disease models and gene correction therapies.

## Acknowledgments

R.M. was funded through NRSA Institutional Predoctoral Training (T32). Grant. R.O.B. was supported through an Individual Postdoctoral grant (DFF–1333-00106B) and a Sapere Aude, Research Talent grant (DFF–1331-00735B), both from the Danish Council for Independent Research, Medical Sciences. MHP thanks the Laurie Kraus Lacob Translational Scholar Fund, the Amon Carter Foundation, the Tad Taube Program in Neurodegenerative Research for their support of this work. DPD thanks the Children’s Health Research Institute at Stanford for their support. VW is supported by a research fellowship from Deutsche Forschungsgemeinschaft (DFG).

## Author Contributions

R.M., K.I. and M.H.P. conceived and designed the experiments and wrote the manuscript. R.M. and K.I. performed the experiments. K.C., T.N., D.D., J.C., R.B., A.L., M.J. V.W. V.S and H.K. all participated in the design and conception of experiments and provided editoral feedback on the manuscript.

## Materials & Methods

### Cell culture

Human ES H9 cells (WiCell) were used for *MYD88* and *HBB* editing, and the iPSC lines described in figure 1c were the following: line 1= 1759, line 2=iLC13-F1, line 3= iAM9-M2, line 4=TkDA3-4, line 5=iSB7-M3. TkDA3-4 iPSCs and iPSCs derived from SCD patient were established from human dermal fibroblasts (Cell Applications Inc) as described previously^16,14^. Cell lines 1759, iLC13-F1, iAM9-M2, and iSB7-M3 were established from human T cells as described previously^17^. Cells were maintained in mTeSR1 (STEMCELL technologies) on feeder free Matrigel (Corning)-coated plate. Subculture was performed every 5-6 days by EDTA method. For 1 day after plating, 10 μM Y-27632 (Tocris) was added to the medium.

### Electroporation and transduction

ESCs or iPSCs (70-80% confluent) were harvested with Accutase (Life Technologies). 150 μg/mL of SpCas9 (Integrated DNA Technologies) and 87.5μg/mL of sgRNA were mixed at 1:3 molar ratio directly, incubated for 10 min at room temperature, then diluted with 20 uL of Opti-MEM (Thermo Fisher Scientific) supplemented with 7.25 mM of ATP and 11.8 mM of MgCl2 containing 500,000 cells. Then nucleofection was performed using 16-well Nucleocuvette Strip with 4D Nucleofector system (Lonza). For plasmid transduction, 1μg of plasmid was added into solution for nucleofection.

The *HBB* and *MYD88* synthetic sgRNAs used were purchased from TriLink BioTechnologies and Synthego, respectively, with chemically modified nucleotides at the three terminal positions at both the 5′ and 3′ ends. Modified nucleotides contained 2′-O-methyl 3′-phosphorothioate and the *HBB* sgRNA from Tri-Link was HPLC-purified. The genomic sgRNA target sequences, with PAM in bold, are: *HBB*: 5′-CTTGCCCCACAGGGCAGTAA**CGG**-3′ and gRNA *MYD88* 5’-*CGCATGTTGAGAGCAGCCAG****GGG****- 3’*. Immediately after electroporation, cells were transferred into one well of a Matrigel coated 24 well plate containing 500 μl of mTeSR with 10 μM Y-27632. AAV6 donor vector at 100k MOI (vector genomes/cell) was added directly to cells after plating and incubated at 37°C. Media was changed 24 hours later and 10 μm Y-27632 was removed after 2 days of transduction.

### Flow cytometry

For measuring targeted editing frequency, cells were harvested. Then cells were washed with PBS containing 1% human albumin and 0.5 mM EDTA. Data was acquired using an Accuri C6 plus flow cytometer (BD biosciences).

### AAV Production

GFP integration donor containing UbC promoter, turboGFP, and bGH polyA targeting the *HBB* and *MYD88* loci as well as the SCD corrective donor, were cloned into pAAV-MCS plasmid (Agilent Technologies) containing AAV2 ITRs. The *HBB* donor creates an insertion of 2.2kb within exon 1 of *HBB* and contains left and right homology arms that are 540bp and 420bp, respectively, and flank the Cas9 cut site. The *MYD88* donor creates an insertion of 2.3kb within exon 1 of *MYD88* and contains left and right homology arms that are 480bp and 450bp, respectively, and flank the Cas9 cut site. The SCD corrective donor contains a total of 2.4 kb AAV vectors were produced as sequence homology surrounding the Glu6Val mutation^7^. described previously with slight modification^18^. Each 15cm^2^ dish of 293FT cells (Life Technologies) was transfected using PEI along with 6μg ITR-containing plasmid and 22μg pDGM6 (gift from D. Russell), carrying AAV6 cap, AAV2 rep, and adenoviral helper genes. 72h post-transfection, cells were harvested, lysed by freeze-thaw cycles, incubated at 37 45min with TurboNuclease at 250 U/mL (Abnova) and purified using an iodixanol density gradient by ultracentrifugation at 237K *g* for 2h at 18°C. AAV6 vectors were extracted from the 60–40% iodixanol interface and exchanged into PBS with 5% sorbitol using an Amicon centrifugal filter 100K MWCO (Millipore Sigma) following the manufacturer’s instructions. Pluronic acid was added to dialyzed vector solution to a final concentration of 0.001%, aliquoted, and stored at −80°C until use. Vectors were titered using qPCR to measure vector genome concentration as described previously^19^.

### Genotyping and sequence analysis

*HBB* and *MYD88 ESC* clones were amplified with the following primers to check for integration: *MYD88*-FW 5’-GACAAGCTCTCTAACTGGAGAATGA-3’, *MYD88*-REV 5’- CCTAAGAATGGTACCAAGGTAGGTC-3’, *HBB*-FW 5’- TAGATGTCCCCAGTTAACCTCCTAT-3’, *HBB*-REV 5’- TTATTAGGCAGAATCCAGATGCTCA-3’. Sickle cell clones edited with the anti-sickling donor (Figure 3a) were amplified with the following primers to check for integration: *HBB*-SCD‐ FW: 5′-GAAGATATGCTTAGAACCGAGG-3′ and *HBB*-SCD-REV: 5′- CCACATGCCCAGTTTCTATTGG-3′. PCR was done with Phusion High-Fidelity Polymerase (Thermo-Scientific). Sequence analysis were performed McLab (South San Francisco, CA, USA) using gel extracted PCR amplicon obtained by Phusion polymerase. Genomic DNA sample was prepared by QuickExtract DNA Extraction Solution (Epicentre Madison) following the manufacturer’s instructions.

Off-target analysis Gene edited ES/iPS cell lines were tested for off-target editing events predicted for each sgRNA by COSMID46 tool (http://crispr.bme.gatech.edu). COSMID is a website tool that considers guide RNA mismatch, insertions and deletions based on the target sequence. *HBB* and SCD‐ iPSC clones were analyzed for off target effects at chromosome 9 (target: CAGCCCCACAGGGCAGTAAGG) and chromosome 17 (target: CTTGCCCACACAGGCCAGTACGG) using the following primers: Chr9-FW (GGAACCATGGGAAGCATGTG), Chr9-REV (CCAGTTTCTAAGAGCGGTGG), Chr17-FW (TGAGCCAAGATTGTGCCATG), and Chr17-REV (AGAGACGGGGAGAAAAGTGT). See supplemental figure 1 for off target results. *MYD88* clones were analyzed for off target effects at chromosome 6 (target: AGCATGCTGAGAGCAGCCAGCGG) and chromosome 1 (target: CGGATGTTGAGAGCAGCCACGGG) using the following primers: Chr6-FW (TGTGTTCTCTTGGTCCTCAGA), Chr6-REV (ACAGAGGGCTTGCCTTCTAA), Chr1-FW (TGTGTTCTCTTGGTCCTCAGA), and Chr1-REV (ACAGAGGGCTTGCCTTCTAA). For off target results in SCD edited iPSCs, refer to figure 1 in supplemental figures.

### Glu6Val allele correction analysis by Nested-ddPCR

SCD-iPSCs edited at *HBB* were harvested 7 days post nucleofection and then analyzed for correction frequencies of the Glu6Val mutation by Nested-ddPCR. An ‘In-Out’ PCR approach (Supplemental Figure 2a) was performed to amplify the *HBB* locus, as to ensure exclusion of episomal AAV6 from analysis. The following primers were used to generate an *HBB*-specific 1.4 kb band: FW(out): 5’-AGGAAGCAGAACTCTGCACTTCA -3’ and REV(in): 5’- AGTCAGTGCCTATCAGAAACCCAAGAG -3’. The PCR product was purified and then diluted to 10 μg/μL in nuclease-free water and then ddPCR was carried out with the PCR product generated as the template. The protocol used a 2-probe set on the sample amplicon, where one probe (HEX) binded the integrated sequence and the other probe (FAM) is a reference sequence downstream of the Cas9 break site. To calculate the percentage of alleles that had integration, we divided the poisson-corrected copies/μL HEX (HR)/FAM (Ref). The following primer/probes were used in the ddPCR reaction: HR probe (HEX)- 5’- TGACTCCTGAGGAAAAATCCGCAGTCA -3’, reference probe (FAM)-5’- ACGTGGATGAAGTTGGTGGTGAGG -3’, nested-forward 5’- TCACTAGCAACCTCAAACAGAC -3’ and nested-rev 5’-CCTGTCTTGTAACCTTGATACC -3’. Duplicates of the same sample were analyzed resulting in the average allele correction of 0.82% for AAV6 only samples and 63.22% for AAV6+RNP samples which was the final result for our edited SCD iPSC experiment.

### Statistical analysis

All data are presented as means ± SD. Error bars are based on SEM. The statistical significance of the observed differences was determined with two-tailed Student’s t tests for pairwise comparisons.

---

**Supplemental Figure 1.**
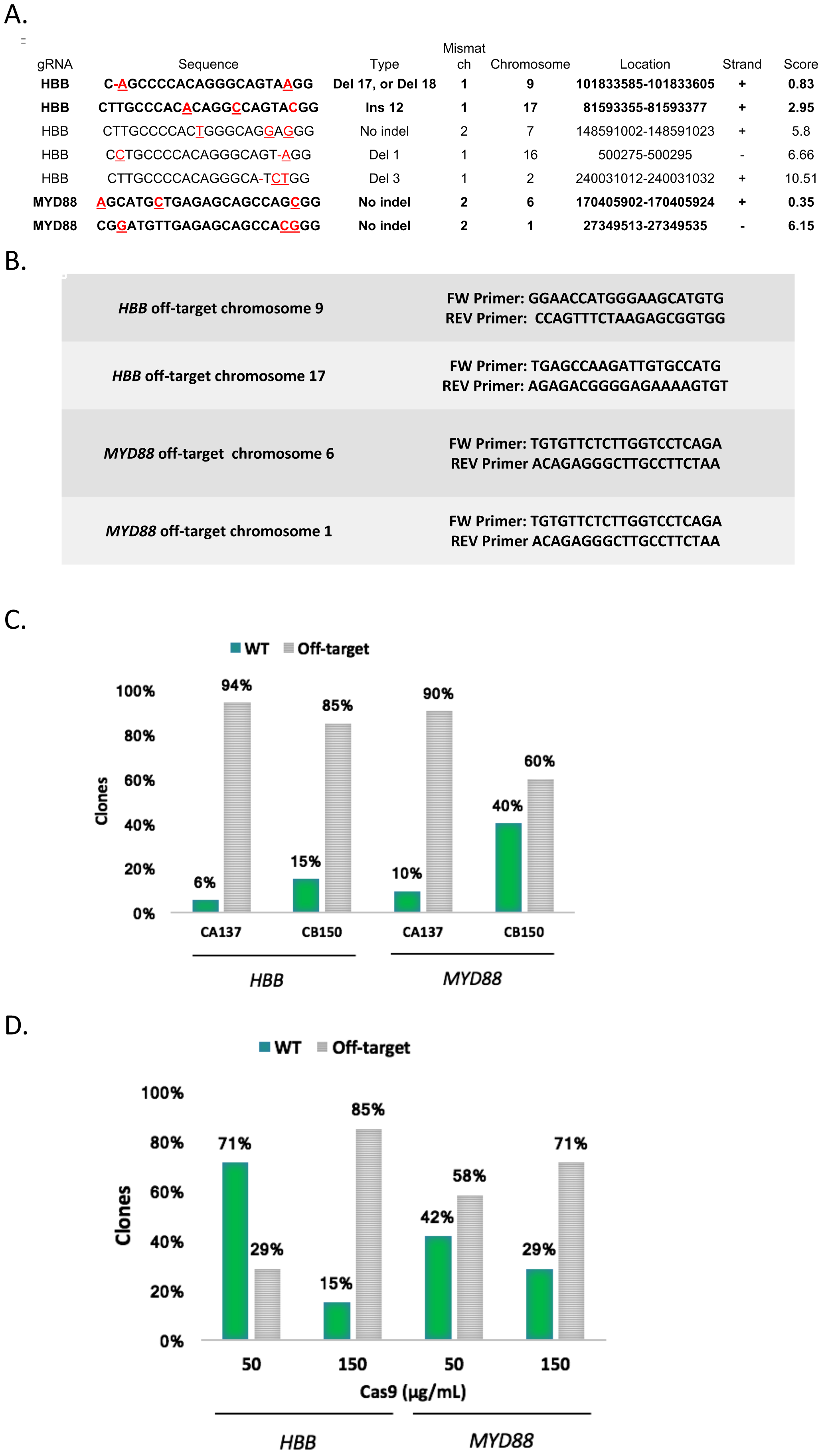
Off-target analysis of edited clones. **a.** Off-target sequences as predicted by COSMID. Top predicted off-target sites are depicted for both *HBB* and *MYD88.* Top two (lowest score) were analyzed. **b.** Primer sequences used to amplify off-target regions. PCR for off-target analysis was done using Phusion Polymerase and gel extracted for sequencing. **c.** Off-target analysis for both *MYD88* and *HBB* clones for CA137 nucleofection protocol and CB150. Both were nucleofected using the optimized Cas9 RNP protocol (150μg). CA137 yields clones with a greater amount of off-target activity. Off-target is considered having indels at one or both of the regions analyzed. WT sequences have no indels at either region. **d.** Off-target analysis for both *MYD88* and *HBB* clones comparing experiments that were done with 50μg and 150 μg of Cas9. All experiments were done with nucleofection code CB150 prior to optimization. Higher Cas9 concentration is correlated with more off-target effects in both *HBB* and *MYD88*.

**Supplemental Figure 2.**
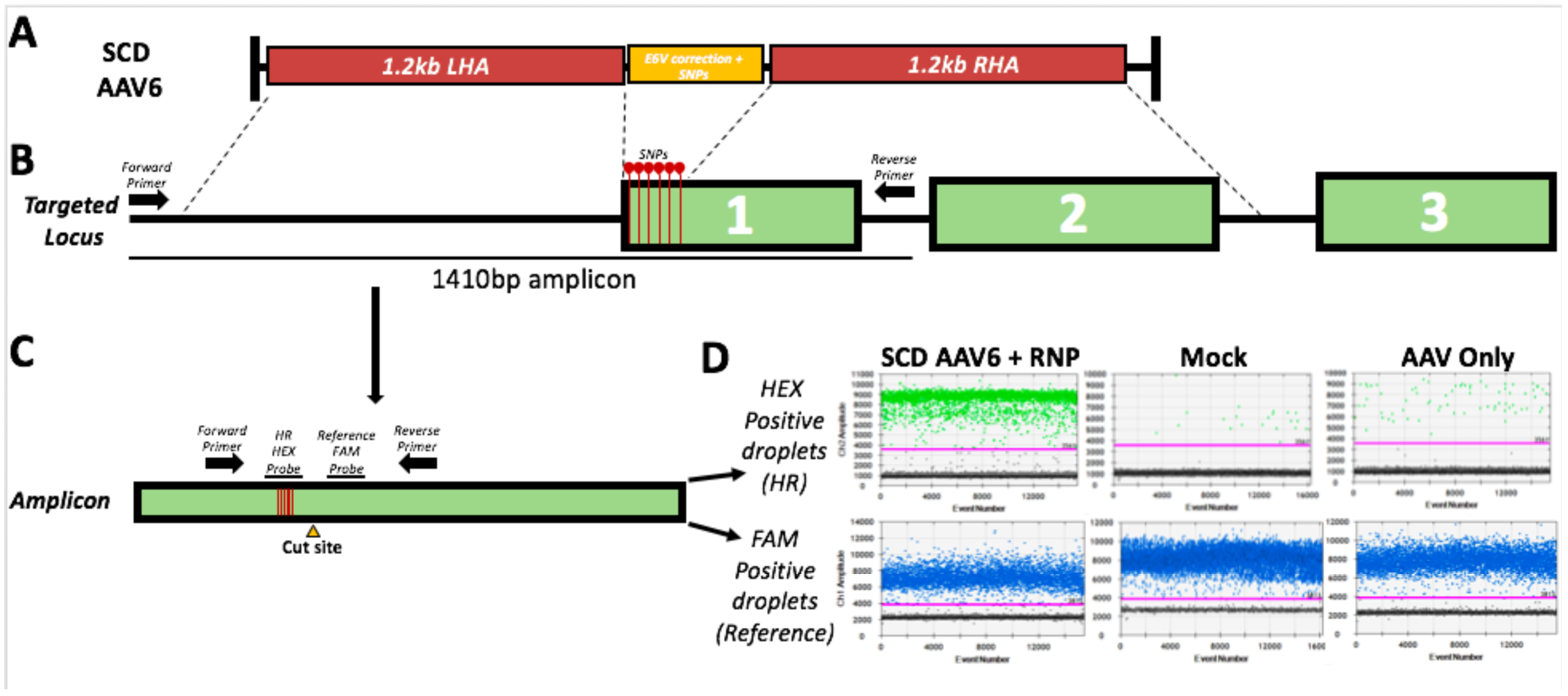
Allele correction analysis by nested-ddPCR after targeting HBB with the E6V allele corrective SCD AAV6 vector. **a.** Schematic of the SCD AAV6 corrective vector and location of homology arms relative to the ‘nested’ primers. **b.** Schematic of the primer binding sites that make the initial HBB-specific 1.4kb amplicon, which excludes episomal AAV6 from analysis. **c.** Schematic of how ddPCR primers/probes bind to PCR product generated from above. A 142 bp amplicon is generated while using a 2-probe set, an HR HEX probe and a reference FAM probe. The HR probe binds to the E6V SNP correction plus the 5 other SNPs introduced by the SCD AAV6 vector and the reference probe binds downstream of the break site to a region native to the HBB region. **d.** Diagrams of the droplet analysis with clear separation shown between positive and negative droplets.

**Supplemental Figure 3.**
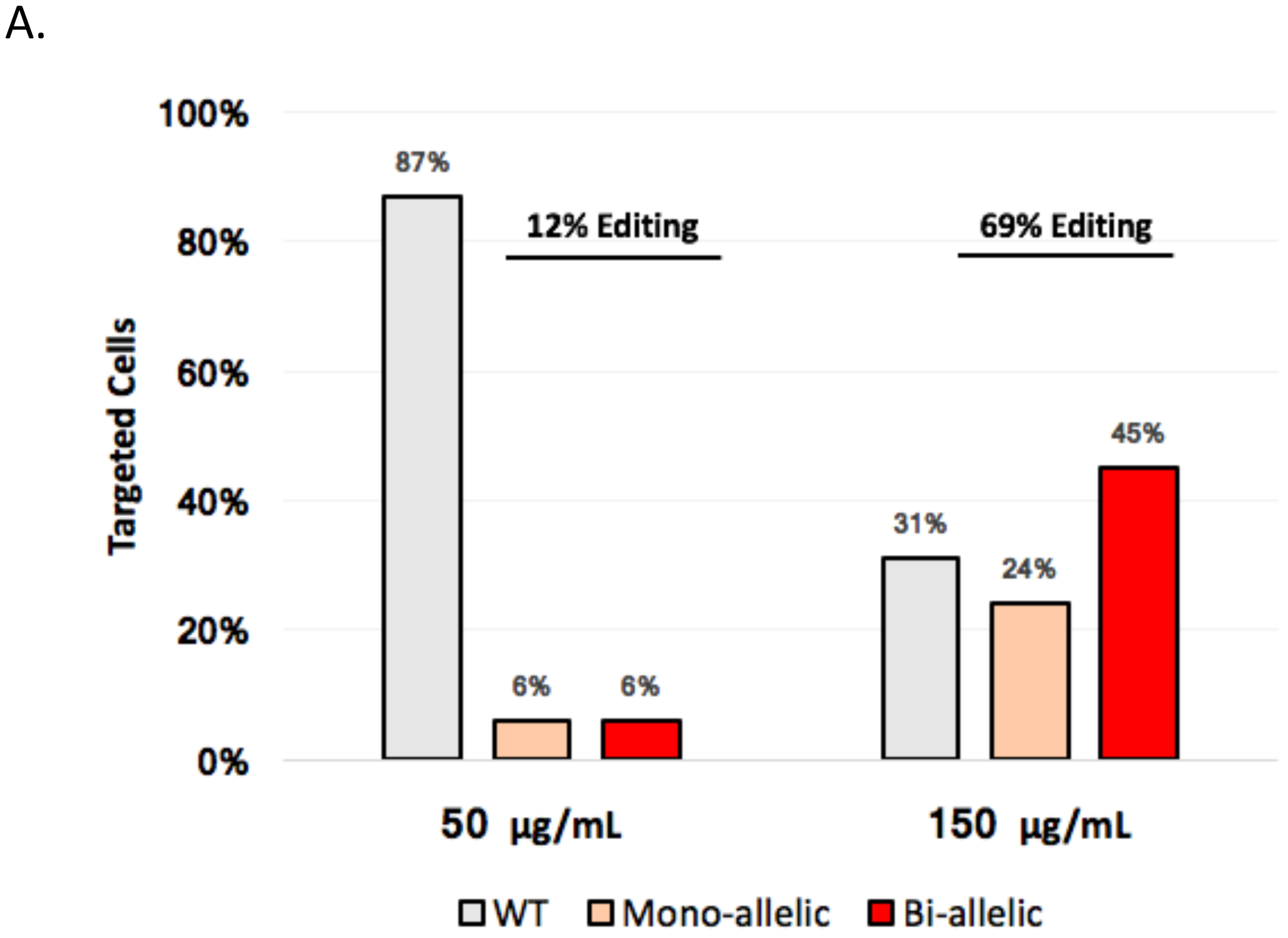
Allele Distribution after Editing using Different Amounts of Cas9/sgRNA RNP. **a.** Distribution of edited cell population after single cell cloning comparing 50μg and 150μg of Cas9. Both experiments were done using CB150 code. Increasing Cas9 not only increases editing frequency, but also increases percentage of bi-allelic integration.

## References

1. Liang, X., Potter, J., Kumar, S., Ravinder, N. & Chesnut, J.D. Enhanced CRISPR/Cas9-mediated precise genome editing by improved design and delivery of gRNA, Cas9 nuclease, and donor DNA. J. Biotechnol. 241, 136–146 (2017).

2. Yang, L. et al. Optimization of scarless human stem cell genome editing. Nucleic Acids Res. 41, 9049–9061 (2013).

3. Wang, G. et al. Efficient, footprint-free human iPSC genome editing by consolidation of Cas9/CRISPR and piggyBac technologies. Nat. Protoc. 12, 88–103 (2017).

4. Zhu, Z., Verma, N., González, F., Shi, Z.-D. & Huangfu, D. A CRISPR/Cas-Mediated Selection-free Knockin Strategy in Human Embryonic Stem Cells. Stem Cell Reports 4, 1103–1111 (2015).

5. Li, X. et al. piggyBac transposase tools for genome engineering. Proc. Natl. Acad. Sci. 110, E2279–E2287 (2013).

6. Yoshimi, K. et al. SsODN-mediated knock-in with CRISPR-Cas for large genomic regions in zygotes. Nat. Commun. 7, 1–10 (2016).

7. Dever, D.P. et al. CRISPR/Cas9 β-globin gene targeting in human haematopoietic stem cells. Nature 539, 384–389 (2016).

8. Bak, R.O. et al. Multiplexed genetic engineering of human hematopoietic stem and progenitor cells using CRISPR/Cas9 and AAV6. Elife 6, 1–19 (2017).

9. Bak, R.O. & Porteus, M.H. CRISPR-Mediated Integration of Large Gene Cassettes Using AAV Donor Vectors. Cell Rep. 20, 750–756 (2017).

10. Hendel, A. et al. Chemically modified guide RNAs enhance CRISPR-Cas genome editing in human primary cells. Nat. Biotechnol. 33, 985–989 (2015).

11. Cradick, T.J., Qiu, P., Lee, C.M., Fine, E.J. & Bao, G. COSMID: A web-based tool for identifying and validating CRISPR/Cas off-target sites. Mol. Ther. - Nucleic Acids 3, (2014).

12. Cradick, T.J., Fine, E.J., Antico, C.J. & Bao, G. CRISPR/Cas9 systems targeting βglobin and CCR5 genes have substantial off-target activity. Nucleic Acids Res. 41, 9584–9592 (2013).

13. DeWitt, M.A. et al. Selection-free genome editing of the sickle mutation in human adult hematopoietic stem/progenitor cells. Sci. Transl. Med. 8, 360ra134–360ra134 (2016).

14. Sebastiano, V. et al. In situ genetic correction of the sickle cell anemia mutation in human induced pluripotent stem cells using engineered zinc finger nucleases. Stem Cells 29, 1717–26 (2011).

15. Kim, S., Bae, T., Hwang, J. & Kim, J.S. Rescue of high-specificity Cas9 variants using sgRNAs with matched 5’ nucleotides. Genome Biol. 18, (2017).

16. Takayama, N. et al. Transient activation of *c-MYC* expression is critical for efficient platelet generation from human induced pluripotent stem cells. J. Exp. Med. 207, 2817–2830 (2010).

17. Nishimura, T. et al. Generation of rejuvenated antigen-specific T cells by reprogramming to pluripotency and redifferentiation. Cell Stem Cell 12, 114–126 (2013).

18. Khan, I.F., Hirata, R.K. & Russell, D.W. AAV-mediated gene targeting methods for human cells. Nat. Protoc. 6, 482–501 (2011).

19. Aurnhammer, C. et al. Universal Real-Time PCR for the Detection and Quantification of Adeno-Associated Virus Serotype 2-Derived Inverted Terminal Repeat Sequences. Hum. Gene Ther. Methods 23, 18–28 (2012).

